# Characterizing CSNK2A1 Mutant-Induced Morphological Phenotypes in Zebrafish (Danio rerio): Insights into Okur-Chung Neurodevelopmental Syndrome (OCNDS)

**DOI:** 10.1101/2024.01.09.574075

**Authors:** Katie Hassett, Sai Srihaas Potu, Aravind Sankaramoorthy, Sukanniya Kaneshamoorthy, Kamawela Leka, Matthew J. Huentelman, Vinodh Narayanan, Sampath Rangasamy

**Affiliations:** Neurogenomics Division, Translational Genomics Research Institute (TGen), Phoenix, Arizona, USA; Center for Rare Childhood Disorders (C4RCD), Translational Genomics Research Institute (TGen), Phoenix, Arizona, USA

**Author notes:** Correspondence: Sampath Rangasamy, or Vinodh Narayanan Center for Rare Childhood Disorders (C4RCD), Neurogenomics Division, Translational Genomics Research Institute (TGen), Phoenix, Arizona, United States.

**Keywords:** Okur-Chung Neurodevelopmental Syndrome, CSNK2A1, Casein Kinase 2 (CK2), Zebrafish phenotype, *Danio rerio*

## Abstract

Okur-Chung Neurodevelopmental Syndrome (OCNDS) is a rare, autosomal dominant disorder caused by heterozygous pathogenic variants in the CSNK2A1 gene. CSNK2A1 encodes the α subunit of protein kinase CK2, involved in diverse biological processes. In 2016, Okur et al. reported the discovery of germline *de novo* missense and canonical splice site mutations in CSNK2A1 in five female patients with OCNDS. The syndrome is characterized by developmental delays, intellectual disability, hypotonia, feeding difficulties, dysmorphic facial features, and disrupted circadian rhythms, leading to sleep disturbances. The complex phenotypic spectrum of OCNDS underscores the need for robust model systems to investigate genotype-phenotype correlations, disease mechanisms, and potential therapies. In this study, we employed an overexpression strategy in a zebrafish model to investigate the functional consequences of select CSNK2A1 variants implicated in OCNDS. Our findings revealed distinct morphological phenotypes resulting from the overexpression of different CSNK2A1 mutants, indicating a direct correlation between genetic alterations and phenotypic manifestations. Notably, the CSNK2A1 p.Arg191Ter (R191X) mutation had a significant impact on the phenotype. Co-injection of wild-type CSNK2A1 mRNA with mutant CSNK2A1 mRNA rescued morphological abnormalities in zebrafish embryos. Overall our study highlights the utility of zebrafish as an adaptable model system for examining the functional impact of CSNK2A1 mutations and exploring novel therapeutic avenues.

## Introduction

Okur-Chung Neurodevelopmental Syndrome (OCNDS) (OMIM# 617062) is an autosomal dominant disorder caused by heterozygous mutations in the CSNK2A1 gene located on chromosome 20p13 (Okur et al., 2016; Wirkner et al., 1994, 1998). CSNK2A1 encodes the α subunit of the highly conserved serine/threonine protein kinase CK2 (OMIM# 115440), a tetrameric protein composed of two α subunits and two β subunits (Chen et al., 2023; Trembley et al., 2023). CK2 is widely expressed in human tissues and regulates multiple biological processes such as cell proliferation, differentiation, development, circadian rhythm, and apoptosis (Halloran et al., 2022). CK2 is an early recognized protein kinase that phosphorylates various substrates with broader specificity ( Wirkner et al., 1994; Wirkner et al.,1998; Meggio & Pinna, 2003; Ruzzene et al., 2011; Borgo et al., 2021). The aberrant functioning of CK2 is associated with a variety of conditions and diseases, such as immune dysfunction, autoimmune diseases, neurodegenerative disorders, including Alzheimer’s and Huntington’s disease, metabolic disorders, and cancer (Abi Nahed et al., 2020; Borgo et al., 2021; Franchin et al., 2017; Lettieri et al., 2019).

In 2016, Okur et al. reported the discovery of germline *de novo* missense and canonical splice site mutations in CSNK2A1 in five female patients with neurodevelopmental syndrome (Okur et al., 2016; Owen et al., 2018). The clinical features commonly observed in individuals with OCNDS include developmental delays, intellectual disability (usually in the mild-to-moderate range), hypotonia, feeding difficulties, dysmorphic facial features, and disrupted circadian rhythms leading to sleep disturbances (Okur et al., 2016; Owen et al., 2018). In addition, various nonspecific clinical features, behavioral problems, speech problems, microcephaly, macrocephaly, short stature, gastrointestinal issues, pachygyria on brain MRI, and dysmorphic features have been frequently described (Okur et al., 2016). Some patients have also reported immune dysfunction, congenital heart defects, and seizures (Chiu et al. 2018; Xu et al. 2020; Wu et al. 2021).

Despite advancements in the understanding of the genetic underpinnings of OCNDS, significant gaps remain in our knowledge of the molecular mechanisms driving this syndrome. Since its initial description by Okur et al. in 2016, the specific genetic mechanisms by which CSNK2A1 missense mutations lead to OCNDS remain elusive. This knowledge gap extends to the explanation of the phenotypic variability observed in individuals with OCNDS, an aspect that is still poorly understood (Wu et al., 2021). The complex phenotypic spectrum of OCNDS underscores the need for the development of robust model systems to bridge the gap between genetic discoveries, genotype-phenotype correlations, molecular mechanisms of disease pathogenesis, and therapeutic advancements.

CSNK2A1, the most abundant subunit of CK2, exhibits a significant degree of evolutionary conservation between species (Graham & Litchfield, 2000). Mice in which CSNK2A1 has been completely knocked out (*Csnk2a1*^−/−^) die during mid-embryogenesis, whereas heterozygous (*Csnk2a1*^+/−^) mice appear to be completely normal (Lou et al., 2008). Disease-causing CSNK2A1 mutations, predominantly characterized by single-nucleotide changes resulting in missense alterations, frameshifts, or premature termination codons, are dispersed throughout the gene with notable hotspots at K198R and R47G (Chung & Okur, 1993). The functional impact of pathogenic CSNK2A1 variants has yet to be comprehensively investigated in animal models, and the development and detailed characterization of mouse models with all known CSNK2A1 mutations pose considerable challenges. This highlights the need for additional animal models, such as zebrafish (*Danio rerio*). Recent breakthroughs have underscored the potential of zebrafish as model organisms, presenting a compelling alternative to conventional rodent models. Owing to their reduced cost, rapid generation time, and high fecundity, zebrafish are particularly suitable for early drug discovery and genetic studies. The zebrafish genome contains CSNK2A1, a human ortholog of the CSNK2A1 protein, which shares 88% identity and analogous functional attributes. This genetic similarity provides a robust foundation for modeling human neurodevelopmental disorders in the zebrafish.

In this study, we explored zebrafish model to study the effects of CSNK2A1 variants that cause OCNDS and elucidate the underlying developmental and behavioral mechanisms. We used an overexpression approach to investigate the functional consequences of disease-causing CSNK2A1 variants in zebrafish embryos. This approach is particularly well suited for analyzing mutations associated with dominant-negative effects. However, we also reasoned that this could be applicable to loss-of-function mutants that retain residual activity. The feasibility of our approach is based on the ubiquitous expression of CSNK2A1 throughout zebrafish development and the rapid integration of the newly synthesized CSNK2A1 subunits into the holoenzyme. This integration allows exogenous mRNA to actively integrate into the CK2 holoenzyme and modulate its function. We microinjected zebrafish embryos with wild-type and mutant CSNK2A1 mRNAs and evaluated the resulting morphological and behavioral effects. Overexpression of mutant proteins led to aberrant phenotypes, with the severity of the effects correlating with the specific mutant mRNA injected. Co-injection with wild-type mRNA successfully rescued the observed phenotypes, confirming a direct link between CSNK2A1 mutants and the observed phenotypes. Collectively, our findings demonstrate the utility of zebrafish as a model system for investigating the effects of CSNK2A1 variants and unraveling the mechanisms underlying OCNDS.

## Methods

### Zebrafish Breeding and Maintenance

The wild-type zebrafish AB strain was obtained from the Zebrafish International Resource Center (ZIRC, Eugene) and used according to the EU Directive 2010/63/EU for animal experiments. The zebrafish were maintained and used according to protocols approved by the local IACUC Committee. They were maintained in tanks containing 2 L of water buffered at 7.5±0.5, 500±200 μ, and 28±1°C with a 14-hour light and 10-hour dark cycle. Adult zebrafish were bred by natural crosses, and embryos were collected and maintained according to established protocols (Kimmel et al. 1995). Collected embryos were grown in a dark incubator at 28.5 °C in E3 embryo medium containing 0.3 g/L NaCL, 0.01 g/L KCl, 0.05 g/L CaCl_2_, 0.08 g/L MgSO_4_, and 0.5 mL/L of methylene blue in sterile water until the desired developmental stage. All embryos were humanely euthanized 72 h post-fertilization (hpf).

### CK2 Inhibitor Study

Zebrafish embryos were exposed to varying concentrations of CX-4945 (Selleck Chemicals LLC, Houston, USA) to determine the optimal dose required to show morphological changes as a result of CK2A inhibition. CX-4945 was dissolved in DMSO to prepare solutions at concentrations of 0.5 μM, 0.75 μM, 1.0 μM, 1.25 μM, and 1.5 μM. These concentrations were utilized to determine the optimal dose for subsequent experiments. Embryos were also exposed to the vehicle control (DMSO) and no treatment control (E3 medium). To elucidate the dose-dependent effects of CX-4945 on zebrafish embryonic development, we selected visibly healthy embryos at 24 hours post-fertilization (hpf) for experimentation. These embryos were then exposed to varying concentrations of CX-4945. Subsequent developmental changes were monitored continuously until 72 hpf at a constant temperature of 28.5°C.

### Microinjections of mRNA

The ready to inject wild-type and human mutant CSNK2A1 ( p.Arg191Ter (R191X), c.571C>T; p.Tyr50Cys (Y50C), c.149A>G; p.Asp156Glu (D156E), c.468T>A; p.Arg47Gln (R47Q), c.139C>G; and p.Lys198Arg (K198R), c.592A>G.) capped mRNA were synthesized by VectorBuilder Inc. (https://en.vectorbuilder.com/resources/vector-system/pT7_mRNA.html). A stock solution of injection medium was prepared with 1 μL of eGFP mRNA, with 1 μL of p53 morpholino (oligo), 0.5 μL of phenol red dye, and the selected and optimized dosage of wild type or mutant CSNK2A1 mRNA (10 pg), and adjusted to a total of 10 μL with ddH_2_O. eGFP mRNA was added to verify efficacy of microinjection at 24 hpf by observing *in vivo* green fluorescent protein (GFP) expression using fluorescence microscopy. Microinjections of the wild-type and mutant mRNA were performed at the single-cell stage of embryonic development. Embryos were loaded into a 2% agarose gel mold and a pre-pulled glass needle (World Precision Instruments, Sarasota) was filled with 10 μL of the injection medium. The injection volume was standardized to ensure consistency across all experimental groups. Post-injection, embryos were incubated at 28.5°C in E3 medium.

### Morphological and Phenotypic Analysis

At 72 hpf, the surviving embryos were anesthetized for morphological examination. Digital images for the morphological analysis of zebrafish embryos were documented using the EVOS FL Cell Imaging System (Thermo Fischer, Waltham). The images were analyzed for phenotypic alterations including yolk sac aberration, tail fin reduction, spinal curvature, and head size. Embryos were categorized into three groups based on the severity of their phenotypic presentation: severe (three or more phenotypes), moderate (two phenotypes), and mild (one phenotype). Embryo survival was monitored at 24, 48, and 72 hpf. Survival rates for the CK2 inhibitor study were calculated as the number of surviving embryos divided by the total number of injected embryos in each group. Survival rates of the microinjected embryos were calculated relative to the control-injected group (number of surviving embryos within the wild-type or mutant groups divided by the number of surviving embryos within the control-injected group).

### Statistical Analysis

The statistical significance of the differences in survival rates and phenotypic severity between the experimental and control groups was determined using the chi-square test for categorical data. Statistical significance was set at p < 0.05. Statistical analyses were conducted using GraphPad Prism 10.

## Results

### Dose Dependent Effect of CK2 inhibitor on Zebrafish Phenotype

CX-4945 is a potent and selective orally bioavailable small-molecule inhibitor of CK2 that is widely used in *in vitro* studies to understand CK2 function. The effect of the CK2 inhibitor was determined using CX-4945 concentration (0.25 μm - 2.5 μm). Initial screening performed to determine the survival rate of embryos showed an increase in embryo mortality at CX-4945 concentrations of 1.25 μM and 1.50 μM (Figure 1A). Morphological analysis of the hatched embryos showed distinct phenotypes, such as an abnormal yolk sac, reduced tail fin size, and curved spines (Figure 1C). CX-4945 concentrations of 0.75 μM - 1.50 μM displayed abnormal phenotype ranging from mild to moderate (Figure 1B). However, concentration 1.0 μM and 1.5 μM displayed significant morphological abnormalities, such as aberrant yolk sac, reduced tail fin size, and curved spine (Figure 1C).

**Figure 1:**
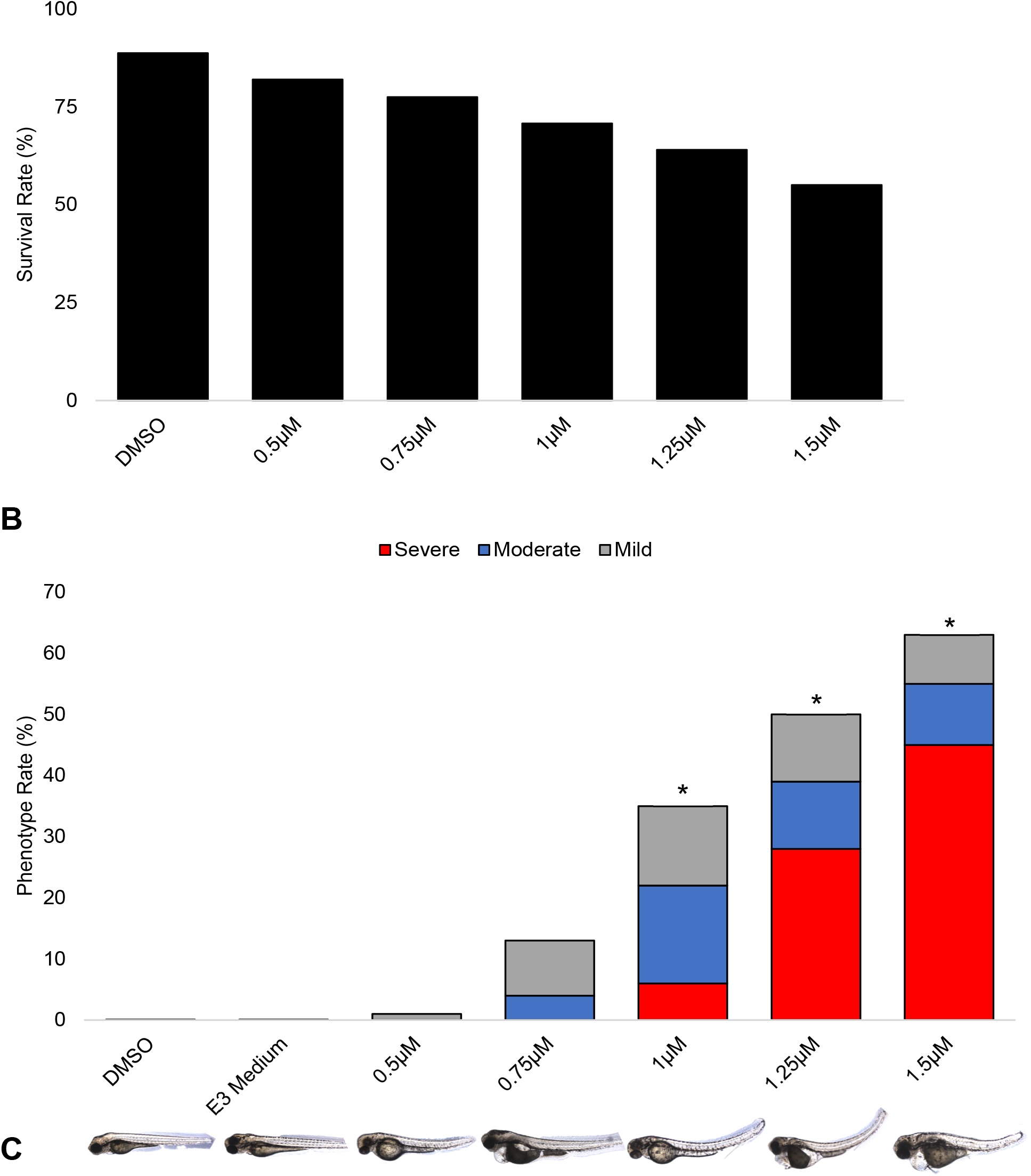
Dose-Dependent Effects of CK2 Inhibitor CX-4945 on Zebrafish Embryos. (A) Survival Rate Analysis: The survival rate of zebrafish embryos treated with varying concentrations of CK2 inhibitor CX-4945. Rates are presented relative to control group. (B) Phenotypic Analysis: Assessment of phenotypic changes in zebrafish embryos at different concentrations of CK2 inhibitor. Severity of phenotypic changes was quantified using a symptom scoring system detailed in the ‘Morphological and Phenotypic Analysis’ section of the Methods. Data represent mean values; error bars indicate standard deviation across five independent trials. Sample size: n > 30 embryos per condition. Statistical significance indicated by * (p<0.05). (C) Morphological Assessment: Representative images depicting the gross morphology of zebrafish embryos following treatment with CX-4945.

### Impact of CSNK2A1 Mutation on Zebrafish Phenotype

To investigate the effect and severity of some common human CSNK2A1 variants (D156E, K198R, R47G, Y50C, and R191X) found in patients, wild-type and mutant CSNK2A1 mRNA carrying those variants were injected into zebrafish embryos at the single-cell stage. The survival and phenotypic outcomes of the injected embryos were compared at 72 hpf in the control-injected group. Our data suggest that embryos injected with the CSNK2A1 variant D156E displayed increased mortality rate (Figure 2A), whereas K198R, R47G, Y50C, and R191X showed a moderate increase in mortality rate compared to the control-injected embryos (Figure 2A). Morphological analysis suggested varying degrees of severity, with more severe morphology observed in variants R191X and Y50C (Figure 2B). Our results indicated that Y50C and R191X mutations had severe and consistent phenotypes, which led to their selection as primary candidates for rescue analysis. The results indicated that the overexpression of disease-causing CSNK2A1 variants resulted in variable phenotypic manifestations (Figure 2C). Notably, the absence of phenotypic deviation in the wild-type mRNA-injected group confirms that the overexpression of wild-type mRNA is not detrimental to embryonic health.

**Figure 2:**
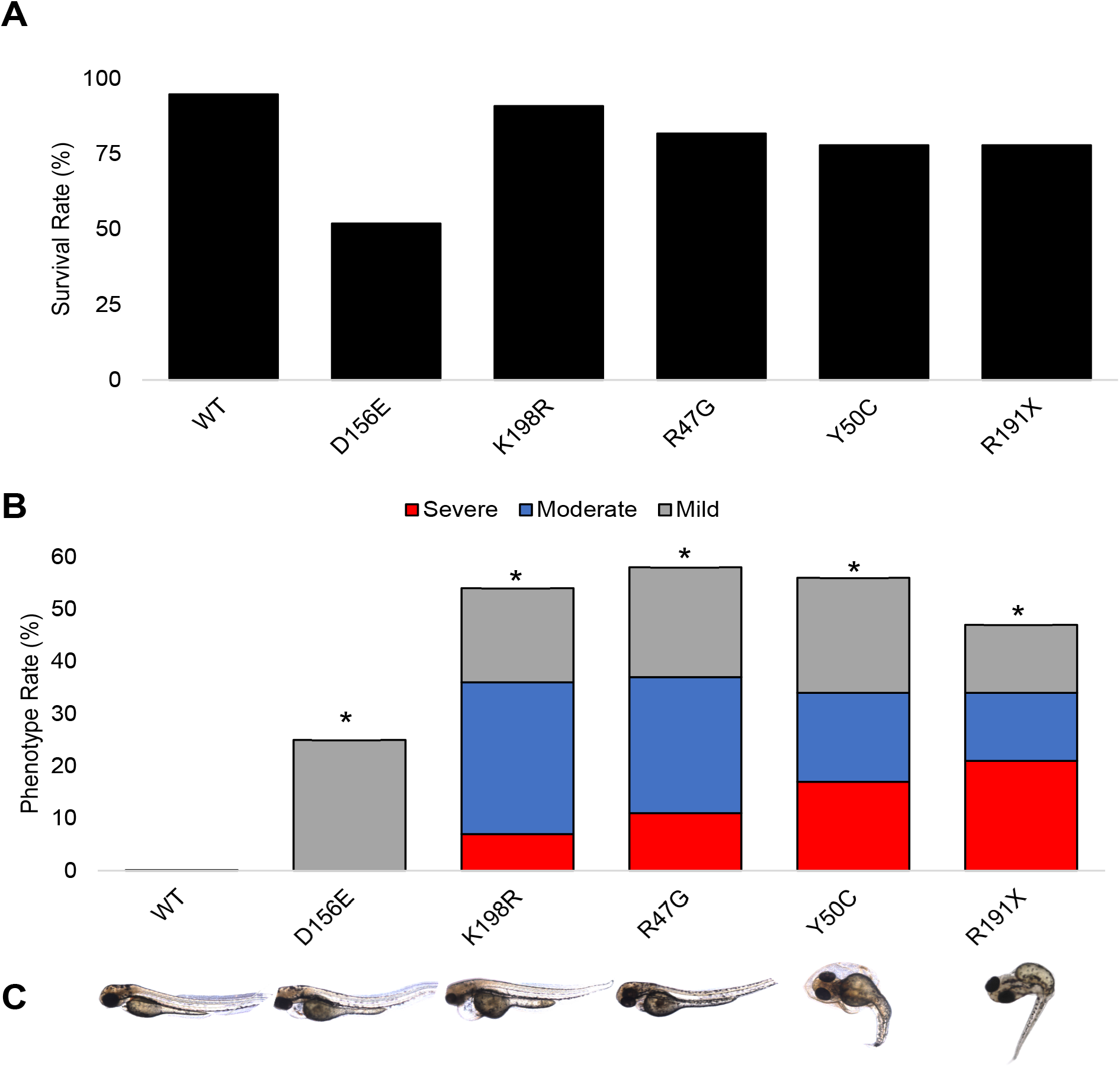
Effects of Overexpression of Wild Type and Mutant CSNK2A1 mRNA. Zebrafish embryos at the single cell stage were injected with a cocktail containing either mutant or wild-type CSNK2A1 mRNA, eGFP mRNA and p53 oligo. Embryos without the expression of eGFP at 24 hpf were excluded to ensure delivery of the cocktail. The survival rate (A) and phenotype severity (B) of injected embryos were assessed relative to control embryos. A) Survival rate of the injected embryos. (B) Phenotypic analysis of the injected zebrafish embryos expressed as percentage severity. n > 30 per condition, *p<0.05. (C) shows the gross morphology of embryos treated with either wild-type or mutant CSNK2A1 mRNA, highlighting key morphological differences.

### Rescue of CSNK2A1 Mutant Phenotype

To evaluate the potential of the wild-type protein to rescue the mutant phenotype, and to elucidate whether the observed developmental defects were attributable to excessive protein expression or a dominant-negative effect of the mutant proteins, co-injection experiments with varying ratios of wild-type and mutant CSNK2A1 mRNA were performed. When an equivalent total quantity of mRNA was maintained, co-injection of the R191X mutant and wild-type mRNA in a 1:2 ratio significantly rescued the phenotype induced by the R191X mutation (Figure 3C & 3E) without significant changes in the survival rate (Figure 3A). Conversely, co-injection of equal amounts of mutant and wild-type CSNK2A1 mRNA resulted in a significant attenuation of the combined phenotypic abnormalities associated with the Y50C variant (Figure 3D & 3F) without significant changes in the survival rate (Figure 3B). These findings imply that the observed developmental defects in zebrafish are not solely caused by the overexpression of mutant proteins but also involve the dominant-negative effects of these aberrant proteins. Supplementary Table 1, offers detailed information on the phenotype of zebrafish embryos from the study.

**Figure 3:**
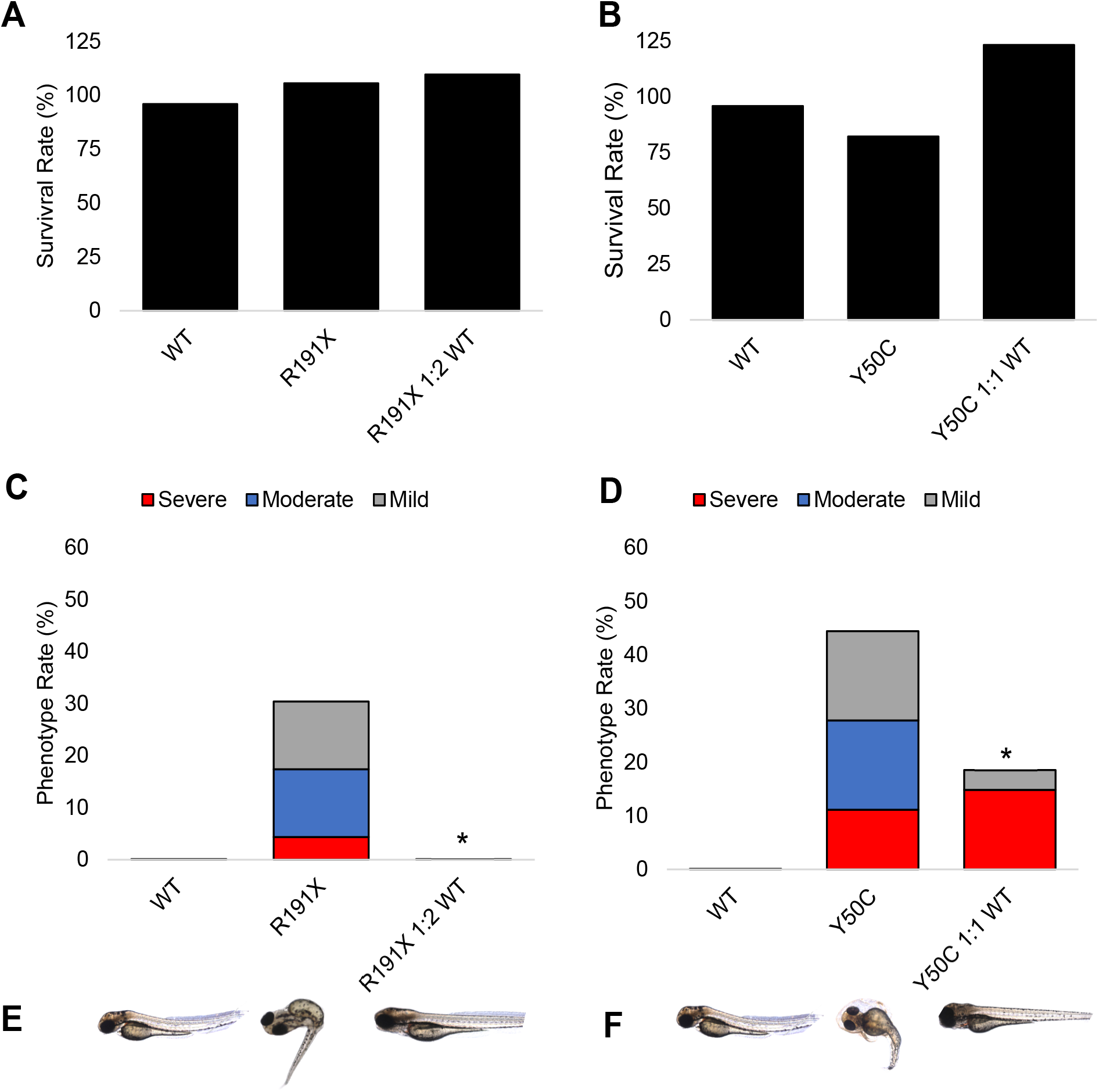
Evaluation of Co-expression of Wild-type and Mutant CSNK2A1 mRNA. Zebrafish embryos were injected with a cocktail of mutant and/or wild-type mRNA, eGFP, and p53 oligo while in the single cell stage. Only embryos exhibiting green fluorescence were included in subsequent analyses. Survival and phenotype rates were assessed relative to control injected embryos. (A and B) Show survival rates, (C and D) Phenotype analysis of the embryos injected with wild-type mRNA, mutant mRNA, and a combination of both wild-type and mutant mRNA. Phenotypic analysis of the injected zebrafish embryos expressed as percentage severity. (C and F) Gross morphology of zebrafish embryos treated with wild-type and mutant (R191X and Y50C) CSNK2A1 mRNA.

## Discussion

Despite significant progress in the understanding of the genetic etiology of OCNDS, the precise molecular mechanisms driving this syndrome remain largely unexplored. To date, more than 30 heterozygous missense CSNK2A1 mutations have been discovered. The functional consequences of these mutations, however, are not entirely clear. Previous studies have indicated that these mutations can lead to varying degrees of reduced kinase activity in cellular models. (Dominguez et al. 2021). Moreover, Caefer et al. (2022) have suggested that these mutations might not necessarily result in a total loss of function, but could potentially alter the specificity towards different substrates. This emerging evidence points towards a more complex disease mechanism than previously thought, where CSNK2A1 mutations might lead to both loss and gain of function in specific contexts.

The development of better animal models of OCNDS has been challenging because of the phenotypic and genotypic heterogeneity observed in this disorder. Zebrafish offer a promising alternative model system for studying OCNDS owing to their high genetic conservation with humans and their suitability for in vivo experimentation. Zebrafish CSNK2A1 shares 88% structural similarity with human CSNK2A1 at both gene and protein levels, suggesting a high degree of functional conservation. In this study, we overexpressed CSNK2A1 variants associated with OCNDS in zebrafish embryos and observed a range of morphological abnormalities, including an aberrant yolk sac, enlarged head, reduced body size, and curved spines. The severity of these defects correlated with the specific CSNK2A1 variant, and the variants R191X and Y50C led to more pronounced abnormalities. These findings suggest that the overexpression of mutant CSNK2A1 disrupts normal embryonic development and contributes to OCNDS pathogenesis.

Additionally, co-injection of wild-type CSNK2A1 mRNA with mutant mRNA rescued the aberrant phenotypes, indicating the dominant-negative effects of the mutant proteins. Observations from this study suggest that mutant CSNK2A1 proteins could interfere with the function of wild-type CSNK2A1 proteins, leading to observed morphological defects. Furthermore, treatment with the CK2 inhibitor CX-4945 mimicked the morphological abnormalities observed in CSNK2A1 mutant embryos. This suggests that CK2 dysfunction, regardless of the CSNK2A1 mutation, plays a significant role in early embryonic development. Overall, our findings demonstrate the feasibility of using zebrafish models to investigate the functional consequences of CSNK2A1 mutations in OCNDS, and highlight the importance of CK2 in early embryonic development.

In this study, we selected five distinct mutations in human CSNK2A1 to model in zebrafish, which are indicated to cause OCNDS, although their potential functional impacts are not yet clear. Upon marked overexpression of the mutant mRNAs, we observed significant morphological defects such as spinal curvature, aberrant yolk extension, enlarged head, and reduced body length. The observed phenotype is not an artifactual consequence of massive expression of an exogenous protein, since the same levels of wild-type protein had no noticeable effects on morphology or survival. Furthermore, the co-expression of wild-type and mutant proteins prevented phenotypic changes, confirming the specificity of the results. Although we were unable to measure protein levels in embryos, wild-type rescue experiment strongly suggests that the mutant CSNK2A1 mRNA causes an aberrant phenotype. The most probable assumption is that a high level of mutant protein competes for interaction with the CK2 regulatory subunit β to form a compromised CK2 enzyme. However, further in-depth biochemical characterization of each mutant protein is warranted to fully elucidate the molecular mechanisms underlying the observed morphological defects. Additionally, more detailed experiments are needed to delineate the precise mechanisms by which mutant proteins disrupt CK2 signaling and contribute to the observed zebrafish phenotype.

In conclusion, our study has demonstrated that overexpression of CSNK2A1 mutants in zebrafish embryos resulted in distinct aberrant phenotypes. Notably, each variant of the CSNK2A1 gene manifested a unique morphological phenotype, indicating a direct correlation between genetic alterations and phenotypic expression. Among the variants studied, the R191X mutation is particularly noteworthy for its severe phenotypic impact. This finding is significant, as it underscores the potential of this variant to serve as a critical marker for understanding the pathophysiology of OCNDS. Importantly, these results offer a compelling rationale for the development of a mouse model that mirrors the R191X mutation, thereby providing a powerful tool for translational research on OCNDS. The use of zebrafish models in this context is invaluable as they offer a highly relevant and adaptable system for probing the functional impact of CSNK2A1 mutations and for the exploration of novel therapeutic avenues. This approach could not only enhance our understanding of OCNDS at the molecular level, but also paves the way for the development of potential treatments.

## Ethics Statement

The research conducted in this study adhered to the highest standards of ethical conduct and complied with all applicable laws and regulations concerning the use of zebrafish as a model organism. All experimental procedures involving zebrafish were designed and performed in accordance with the principles and guidelines of the Institutional Animal Care and Use Committee (IACUC) (IACUC Protocol# 22153/TGEN).

## Author Contributions

KH, VN and SR contributed to the design and implementation of the research. KH, SP, AS, KL, MH, VN and SR contribute to the analysis of the results and the writing of the manuscript.

## Acknowledgements

This study was supported by private donor funding from the CSNK2A1 Foundation to Dr. Narayanan.

## References

Abi Nahed, R., Reynaud, D., Lemaitre, N., Lartigue, S., Roelants, C., Vaiman, D., Benharouga, M., Cochet, C., Filhol, O., & Alfaidy, N. (2020). Protein kinase CK2 contributes to placental development: physiological and pathological implications. Journal of Molecular Medicine, 98(1), 123–133. 10.1007/s00109-019-01855-0

Borgo, C., D’Amore, C., Sarno, S., Salvi, M., & Ruzzene, M. (2021). Protein kinase CK2: a potential therapeutic target for diverse human diseases. Signal Transduction and Targeted Therapy, 6(1), 183. 10.1038/s41392-021-00567-7

Caefer, D. M., Phan, N. Q., Liddle, J. C., Balsbaugh, J. L., O’Shea, J. P., Tzingounis, A. V., & Schwartz, D. (2022). The Okur-Chung Neurodevelopmental Syndrome Mutation CK2K198R Leads to a Rewiring of Kinase Specificity. Frontiers in Molecular Biosciences, 9, 850661. 10.3389/fmolb.2022.850661

Chen, Y., Wang, Y., Wang, J., Zhou, Z., Cao, S., & Zhang, J. (2023). Strategies of targeting CK2 in drug discovery: challenges, opportunities, and emerging prospects. Journal of Medicinal Chemistry, 66(4), 2257–2281. 10.1021/acs.jmedchem.2c01523

Chiu, A. T. G., Pei, S. L. C., Mak, C. C. Y., Leung, G. K. C., Yu, M. H. C., Lee, S. L., Vreeburg, M., Pfundt, R., van der Burgt, I., Kleefstra, T., Frederic, T. M. T., Nambot, S., Faivre, L., Bruel, A. L., Rossi, M., Isidor, B., Küry, S., Cogne, B., Besnard, T., …Chung, B. H. Y. (2018). Okur-Chung neurodevelopmental syndrome: Eight additional cases with implications on phenotype and genotype expansion. Clinical Genetics, 93(4), 880–890. 10.1111/cge.13196

Chung, W., & Okur, V. (1993). Okur-Chung Neurodevelopmental Syndrome. In M. P. Adam, G. M. Mirzaa, R. A. Pagon, S. E. Wallace, L. J. Bean, K. W. Gripp, & A. Amemiya (Eds.), GeneReviews®. University of Washington, Seattle.

Dominguez, I., Cruz-Gamero, J. M., Corasolla, V., Dacher, N., Rangasamy, S., Urbani, A., Narayanan, V., & Rebholz, H. (2021). Okur-Chung neurodevelopmental syndrome-linked CK2α variants have reduced kinase activity. Human Genetics, 140(7), 1077–1096. 10.1007/s00439-021-02280-5

Franchin, C., Borgo, C., Zaramella, S., Cesaro, L., Arrigoni, G., Salvi, M., & Pinna, L. A. (2017). Exploring the CK2 paradox: restless, dangerous, dispensable. Pharmaceuticals (Basel, Switzerland), 10(1). 10.3390/ph10010011

Graham, K. C., & Litchfield, D. W. (2000). The regulatory beta subunit of protein kinase CK2 mediates formation of tetrameric CK2 complexes. The Journal of Biological Chemistry, 275(7), 5003–5010. 10.1074/jbc.275.7.5003

Halloran, D., Pandit, V., & Nohe, A. (2022). The role of protein kinase CK2 in development and disease progression: A critical review. Journal of Developmental Biology, 10(3). 10.3390/jdb10030031

Lettieri, A., Borgo, C., Zanieri, L., D’Amore, C., Oleari, R., Paganoni, A., Pinna, L. A., Cariboni, A., & Salvi, M. (2019). Protein kinase CK2 subunits differentially perturb the adhesion and migration of GN11 cells: A model of immature migrating neurons. International Journal of Molecular Sciences, 20(23). 10.3390/ijms20235951

Lou, D. Y., Dominguez, I., Toselli, P., Landesman-Bollag, E., O’Brien, C., & Seldin, D. C. (2008). The alpha catalytic subunit of protein kinase CK2 is required for mouse embryonic development. Molecular and Cellular Biology, 28(1), 131–139. 10.1128/MCB.01119-07

Meggio, F., & Pinna, L. A. (2003). One-thousand-and-one substrates of protein kinase CK2? The FASEB Journal, 17(3), 349–368. 10.1096/fj.02-0473rev

Okur, V., Cho, M. T., Henderson, L., Retterer, K., Schneider, M., Sattler, S., Niyazov, D., Azage, M., Smith, S., Picker, J., Lincoln, S., Tarnopolsky, M., Brady, L., Bjornsson, H. T., Applegate, C., Dameron, A., Willaert, R., Baskin, B., Juusola, J., & Chung, W. K. (2016). De novo mutations in CSNK2A1 are associated with neurodevelopmental abnormalities and dysmorphic features. Human Genetics, 135(7), 699–705. 10.1007/s00439-016-1661-y

Owen, C. I., Bowden, R., Parker, M. J., Patterson, J., Patterson, J., Price, S., Sarkar, A., Castle, B., Deshpande, C., Splitt, M., Ghali, N., Dean, J., Green, A. J., Crosby, C., Deciphering Developmental Disorders Study, & Tatton-Brown, K. (2018). Extending the phenotype associated with the CSNK2A1-related Okur-Chung syndrome-A clinical study of 11 individuals. American Journal of Medical Genetics. Part A, 176(5), 1108–1114. 10.1002/ajmg.a.38610

Ruzzene, M., Tosoni, K., Zanin, S., Cesaro, L., & Pinna, L. A. (2011). Protein kinase CK2 accumulation in “oncophilic” cells: causes and effects. Molecular and Cellular Biochemistry, 356(1–2), 5–10. 10.1007/s11010-011-0959-2

Trembley, J. H., Kren, B. T., Afzal, M., Scaria, G. A., Klein, M. A., & Ahmed, K. (2023). Protein kinase CK2 - diverse roles in cancer cell biology and therapeutic promise. Molecular and Cellular Biochemistry, 478(4), 899–926. 10.1007/s11010-022-04558-2

Wirkner, U., Voss, H., Ansorge, W., & Pyerin, W. (1998). Genomic organization and promoter identification of the human protein kinase CK2 catalytic subunit alpha (CSNK2A1). Genomics, 48(1), 71–78. 10.1006/geno.1997.5136

Wirkner, U., Voss, H., Lichter, P., Ansorge, W., & Pyerin, W. (1994). The human gene (CSNK2A1) coding for the casein kinase II subunit alpha is located on chromosome 20 and contains tandemly arranged Alu repeats. Genomics, 19(2), 257–265. 10.1006/geno.1994.1056

Wu, R.-H., Tang, W.-T., Qiu, K.-Y., Li, X.-J., Tang, D.-X., Meng, Z., & He, Z.-W. (2021). Identification of novel CSNK2A1 variants and the genotype-phenotype relationship in patients with Okur-Chung neurodevelopmental syndrome: a case report and systematic literature review. The Journal of International Medical Research, 49(5), 3000605211017063. 10.1177/03000605211017063

Xu, S., Lian, Q., Wu, J., Li, L., & Song, J. (2020). Dual molecular diagnosis of tricho-rhino-phalangeal syndrome type I and Okur-Chung neurodevelopmental syndrome in one Chinese patient: a case report. BMC Medical Genetics, 21(1), 158. 10.1186/s12881-020-01096-w

