## Supplement Table 1 for "Characterizing CSNK2A1 Mutant-Induced Morphological Phenotypes in Zebrafish (Danio rerio): Insights into Okur-Chung Neurodevelopmental Syndrome (OCNDS)"

**Supplemental Table 1:** Zebrafish Phenotype Embryos Following Treatment with CK2 Inhibitor and CSNK2A1 mRNA.

1A) Quantitative analysis of phenotypes in embryos treated with CK2 Inhibitor CX-4945.

1B) Phenotypic outcomes in embryos injected with either wild-type or mutant CSNK2A1 mRNA. 1C) and 1D) Analysis of phenotypic rescue in embryos co-injected with wild type and mutant (R191X and Y50C) CSNK2A1 mRNA. For each category, phenotypic data are presented as percentages.

**1A**

| **Phenotype** | **DMSO** | **Egg Water** | **0.5 µM** | **0.75 µM** | **1 µM** | **1.25 µM** | **1.5 µM** |
| --- | --- | --- | --- | --- | --- | --- | --- |
| **Mild** | 0 | 0 | 1 | 9 | 13 | 11 | 8 |
| **Moderate** | 0 | 0 | 0 | 4 | 16 | 11 | 10 |
| **Severe** | 0 | 0 | 0 | 0 | 6 | 28 | 45 |

**1B**

| **Phenotype** | **WT** | **D156E** | **K198R** | **R47G** | **Y50C** | **R191X** |
| --- | --- | --- | --- | --- | --- | --- |
| **Mild** | 0 | 25 | 18 | 21 | 22 | 13 |
| **Moderate** | 0 | 0 | 29 | 26 | 17 | 13 |
| **Severe** | 0 | 0 | 7 | 11 | 17 | 21 |

**1C**

| **Phenotype** | **WT** | **R191X** | **R191X 1:2 WT** |
| --- | --- | --- | --- |
| **Mild** | 0 | 13 | 0 |
| **Moderate** | 0 | 13 | 0 |
| **Severe** | 0 | 4 | 0 |

**1D**

| **Phenotype** | **WT** | **Y50C** | **Y50C 1:1 WT** |
| --- | --- | --- | --- |
| **Mild** | 0 | 17 | 4 |
| **Moderate** | 0 | 17 | 0 |
| **Severe** | 0 | 11 | 15 |
